# ClinCNV: novel method for allele-specific somatic copy-number alterations detection

**DOI:** 10.1101/837971

**Authors:** German Demidov, Stephan Ossowski

## Abstract

**Motivation:** Large somatic copy number alterations (CNA), short indels and single nucleotide variants (SNVs) are playing important role in cancer development and can serve as a predictor for targeted therapy selection as well as prognostic factor. Genomic microarrays, FISH, MLPA and many other technologies are widely used for detection of CNAs. Whole-genome sequencing (WGS), whole-exome sequencing (WES) and targeted panel sequencing (TPS) are well established, highly accurate tools for detection of SNVs and small indels, but detection of larger structural variants using WGS, WES and TPS data remains challenging. We developed a tool for high-resolution allele-specific detection of somatic CNAs in NGS data using statistical approach.

**Results:** We have developed a new method for read-depth and B-allele frequency (BAF) based multi-sample detection of copy-number changes in paired normal-tumor NGS data and showed its performance using large cohorts of WES and TPS sequenced samples.

**Availability:** ClinCNV is freely available on https://github.com/imgag/ClinCNV.

## I. INTRODUCTION

During the recent decade, big progress in cancer genomics was made. Hundreds of thousands of tumor-normal pairs were sequenced using WGS, WES or TPS approaches for research or clinical purposes. Structural variants play a significant role in cancer development and treatment, so detection of SVs becomes essential for tumor genome analysis. Several tools that successfully decipher a whole variety of structural variants including CNAs in high-coverage WGS data were developed (such as MANTA [Chen et al., 2015]), but CNAs detection in WES and TPS data is still challenging. Nonetheless, targeted sequencing is used more often than WGS in routine clinical practice. The main reasons for this are lower price allowing for deeper sequencing and the possibility to accurately infer alterations within specific cancer-related genomic regions due to high short read coverage in targeted (which usually means important) regions.

We first assembled a set of requirements for CNA detection tools used for clinical diagnostics: 1) ability to work with heterogeneous tumors, 2) high sensitivity even for comparatively short CNAs or sub-clonal events with the low CCF, 3) high specificity even for noisy samples; 4) ability to perform allele-specific calling for the deciphering of complex CNAs, 5) ability to work with all the types of NGS data used for the analysis of cancer genomes, 6) intuitive output visualization for clinical interpretation by clinicians. Several tools have already proven their comparatively good performance in targeted sequencing NGS data, such as AscatNGS [Raine et al., 2016], CNV-Kit [Talevich et al., 2016] and FACETS [Shen et al., 2016], among others, but none of them meets all established requirements perfectly.

To address this issue, we have developed ClinCNV, a tool that utilizes on- and off-target read depth and B-allele frequency signatures to generate genome-wide allele-specific calls. Since the price of sequencing was dropping rapidly during last years, we aimed at the detection of variants in large (>30 samples) cohorts where large parts of the genome (hundreds of genes or more) were sequenced. We analyzed 473 sample pairs from the CLL study ([Puente et al., 2015]) and compared the performance of ClinCNV to other tools as well as to matched microarray data. Additionally, we analyzed 251 clinical samples, sequenced with TPS, and showed the good concordance between our results and the results obtained with the alternative method.

## II METHODS

### i. Input data and normalization

ClinCNV’s normalization flowchart is provided in figure 1. We describe the algorithm in a step-wise manner, moving from the top of the flowchart.

**Figure 1:**
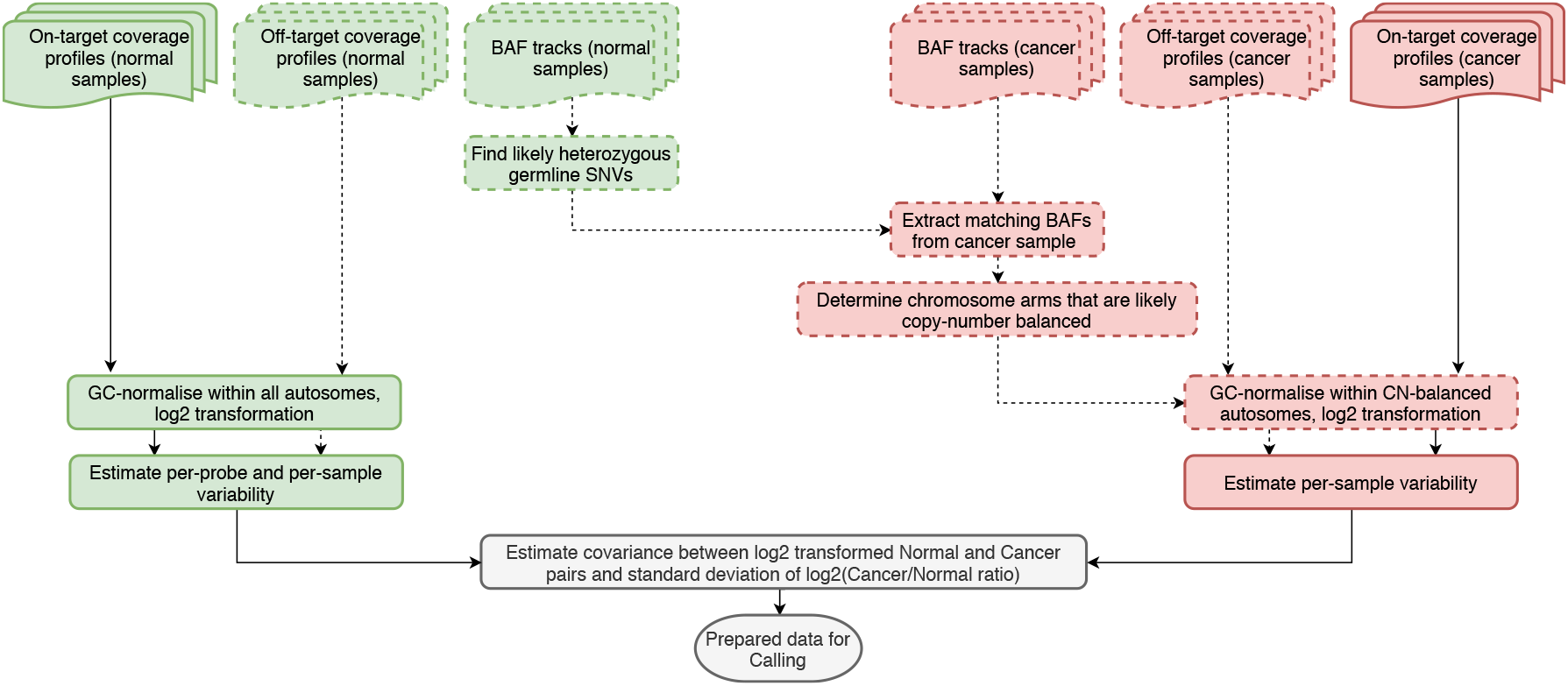
Input data and normalization in Somatic mode of ClinCNV. Dashed line blocks are optional.

#### i.1 Generation of input data for ClinCNV

ClinCNV is a multi-sample caller so all the samples sequenced with the same panel should be analyzed jointly. On-target, off-target read coverage depth is used together with the signal from heterozygous germline SNVs and their matching tumor variants allele frequencies (BAFs). As input, coverage profiles and BAF tracks should be pre-calculated from .BAM and. BED files. To generate BAF tracks, germline SNVs have to be called, or existing databases of populational SNPs may be used. We follow the first approach since it provides more information but may lead to privacy issues, so we intentionally store BAF data (which can be removed immediately after calling) separately from calling results and other input data to prevent accidental revealing of the private variant information.

#### i.2 Exclusion of copy-number imbalanced chromosome arms from further normalization process

In order to facilitate in-depth analysis of allelic imbalances, we use bi-allelic single nucleotide variants, namely their BAFs (ratio between the number of reads supporting the alternative allele of an SNV to the overall coverage of the position). We use only well-covered in both tumor and normal samples SNVs which are likely to be heterozygous in normal tissue.

For tumor samples deeply affected by copy-number alterations, we need to perform all the normalization steps within the genomic regions where the copy number state is expected to be neutral. Ideally we would perform such normalization fo diploid genomic regions only, but we can not infer which regions are diploid and only small part of the genome may remain diploid, thus, we include all copy-number balanced regions and rely on the fact that one copy-number (2 for diploid or 4 for tetraploid tumors) is usually dominating the other copy-number balanced states. We use B-allele frequency as an indicator if the genomic region can be used for normalization or not. Namely, we select chromosome arms which are likely to be copy-number neutral and use only them for normalization.

We test all the SNVs within the chromosome arm and then check the percentage of SNVs that are likely to be different between normal and tumor tissues. We expect such percentage to be high for the chromosome arms highly affected by CNAs (and thus not suitable for the normalization). For each SNV from the chromosome arm, we use the proportion test to determine if the allele ratio for the particular SNV is different between the tumor and healthy tissues. Thenwe count the number of p-values that are below the 0.01 significance threshold and exclude any chromosome where the fraction of significant p-values is bigger than 5% (thus, allowing up to 5% of outlying values). Otherwise, we assume that copy-number change was relatively small or affected only a small part of the chromosome arm, and we include coverages from this arm into the normalization procedure. However, we need at least 5 chromosome arms for performing normalization since variance increases rapidly when using a smaller amount of sequenced genomic material for normalization. Thus, even if no chromosome arms pass the filter, we choose at least 5 chromosome arms with the smallest number of significant p-values.

#### i.3 Normalization of coverage profiles

We perform targeted regions length-based and GC-normalization for tumor and normal samples separately, using only chromosome arms selected at the previous step for tumor sample normalization. log_2_-transformation of ratios is applied for tumor-normal coverage ratios.

#### i.4 Estimation of per-sample and per-region variance of normalized coverage depth

This step is performed in the same manner for off-target and on-target read coverage counts. These two sources of coverage signal have different properties, so we can not normalize them jointly.

To achieve the maximum power of detection, we separate sources of variability, namely – we estimate variances separately for the normal sample and the tumor sample and also we estimate each individual region variance. Since tumor and normal samples’ coverage depths profiles are often highly correlated, we estimate the correlations between them and then infer variance of the log-ratios for each sample pair and each genomic region. Existing tools for CNA detection usually use the assumption that log-ratios of normalized read counts between the same genomic region in tumor and matched normal sample (denoted as *T* and *N*, respectively) follow the Normal distribution.

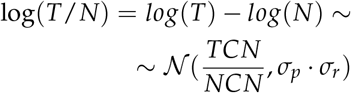

where *TCN* denotes tumor copy number, *NCN* denotes normal copy-number, and 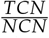 is a subject of CNA detection problem which is possible to solve only knowing log-ratio variance (*σ*_*p*_ denotes the sample pair variance, *σ*_*r*_ denotes the specific individual region variance of log-ratios). Using the normality of log-ratios assumption, we also assume that *T* and *N* follow the log-normal distribution which may be an inaccurate, but useful approximation. Thus, in order to exclude per-sample variability, we standardize log(*N*) and log(*T*) with z-transformation using their per-sample variances estimated directly from the data and median of log(*N*) or log(*T*), respectively:

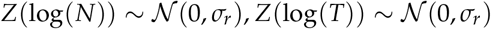

For the tumor sample’s variance we estimate it for each chromosome arm separately and then choose the median value, for normal sample the variance is calculated within all autosomal regions.

Having *Z*(log(*N*)) ~ *Normal*(0, *σ*_*r*_), we can estimate the individual genomic regions coverage depths’ variances from the data. Here and below we always use robust measures of standard deviation (such as *Q*_*n*_, [Rousseeuw et al., 1993]) for estimation. Then we calculate robust correlations between standardized Tumor and Normal normalised counts ([Pasman et al., 1987]). Again, we calculate such correlations for each chromosome arm separately and then choose the median value as a final estimation. In the end, in order to estimate the final variance of log(*T*/*N*), which is equal to 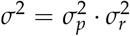, we use the formula:

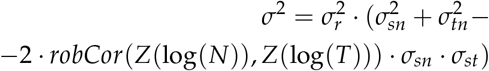

where *σ*_*sn*_ denotes the variance of a normal sample, *σ*_*st*_ denotes the variance of a tumor sample.

In order to estimate variances of regions from X and Y chromosomes, we randomly re-sample log-transformed normalized counts for all males with the counts from females (for chrX) and vice versa for the Y chromosome.

#### i.5 Statistical modeling of BAF

We model BAFs with Normal distribution approximation with an expected value equal to *mappingBias* (*A*/*D*) or *mappingBias* (*B*/*D*), where *A* denotes the number of reads supporting alternative allele of an SNV and *B* denotes the reference allele count, *D* = *A* + *B*. We do not allow the expected value to be below 0.01 or above 0.99. The expected variance is inferred as the Binomial distribution variance. We correct for positive reference alignment bias, estimated as median of BAFs divided by 0.5.

### ii. Calling

Having data normalized and parameters estimated we start calling procedure (fig. 2). At first, we define a set of potential copy-number changes that may occur in a tumor sample at each allele. In the beginning, we investigate major and minor alleles under assumptions of:

1. Discrete set of the potential clonal fraction (starting from 5 to 100% with the step of 2.5%);
2. Minor copy number from 0 to 4;
3. Major copy number from 0 to 30: major and minor copies together are limited to 30 copies.

**Figure 2:**
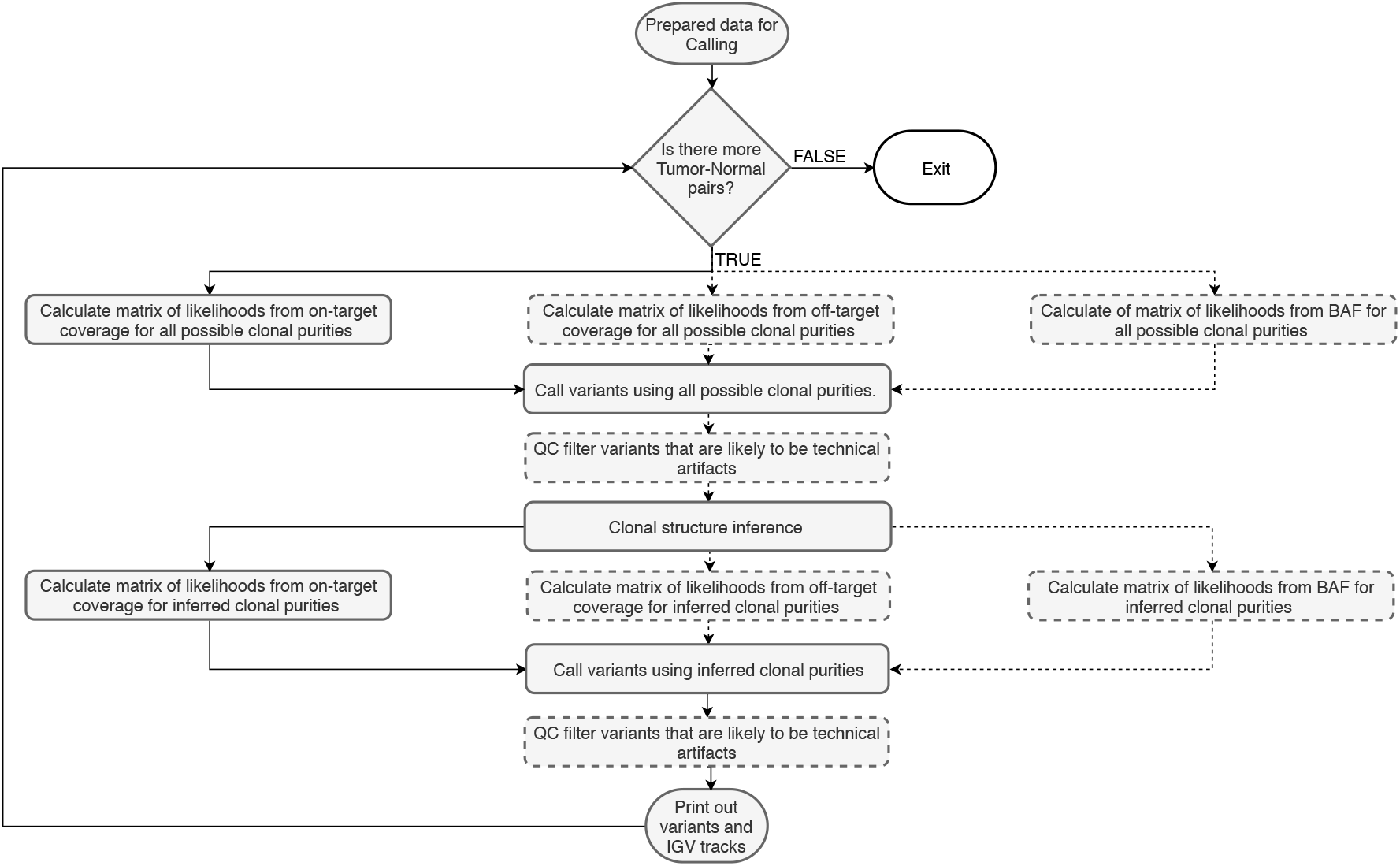
Flowchart of CNA calling. Dashed line blocks are optional.

Having probabilistic models and set of states *S* we can calculate likelihoods of each data point *x*_1_, …, *x*_*n*_ under all these models: 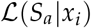. We fill a matrix of size |*S*| × |*G*|, where |*S*| denotes the number of states and |*G*| is the length of the genome with the corresponding likelihoods. It is usually possible to define a baseline copy-number state: it is a diploid state for human or mice autosomes, or a haploid state is a baseline for males’ sex chromosomes. We denote the probabilistic model corresponding to such a baseline state as *S*_*b*_. The problem of finding one CNV in such a matrix may be formulated as:

#### Problem 1

Having matrix of likelihoods of data-points under different states and baseilne state S_b_, identify a pair of indices i, j and state S_a_ ≠ S_b_ such as 1 ≤ i ≤ j ≤ |G| and

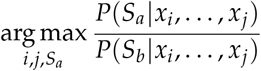

We take logarithms of likelihoods so we will be able to sum them instead of multiplying, so now we have a matrix of 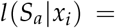 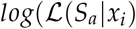. Then, for each genomic region *i* we can switch to likelihood ratio 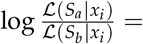 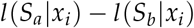. To find segmentsL with the largest sums for one particular state, we can use a well-known maximum subarray sum algorithm ([Bentley et al., 1984]) that solves the following problem in linear time:

#### Problem 2

*Giving a one-dimensional array of numbers A*, 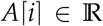, *find indices i and j, 1 ≤ i ≤ j ≤ n, such as* 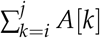 *is a large as possible*.

Whenever we have different signals (such as read depth and B-allele frequency), we can simply sum up matrices of likelihoods from different signal types and perform the same procedure, so it does not affect computational time except for the additional calculations of likelihoods. The actual copy-number state (as well as cancer cell fraction) of the variant is inferred by determining which *S*_*a*_ is the best for the explanation of the observed data comparing to the baseline *S*_*b*_.

Segmentation of a genomic piece into candidate CNV regions is done analogously to CBS. We find one piece that shows the presence of an alternative model at one step of our algorithm, and then we divide the initial genomic piece into three segments: one to the left, one to the right from the discovered segment and the discovered segment itself. We stop segmenting once the next detected segment fails to reach a significance threshold. When the detected variant is shorter than the pre-specified length threshold, but significant, we correct the corresponding likelihood of this potential outlier.

To correct for potential ploidy change we not only select BAF-balanced chromosomes’ arms for normalization, but also try to separate tetraploid regions from diploid, using the assumption that the smallest possible BAF-balanced chromosome arm copy-number is 2 (since, as mentioned in [Shen et al., 2016], long stretches of homozygous deletions most probably lead to cell death and thus are highly unlikely). In order to do this, we sub-select all chromosome arms with small deviations in BAF and calculate coverage baselines (medians of log-ratios of coverages) for all such arms. Then we iteratively merge all arms with differences in medians of their normalized coverage less than 2.5% and choose the smallest merged value that includes at least 10% of markers from BAF-balanced chromosome arms as the baseline.

After we have finished the first round of calling using all possible purities, we calculate the likely sub-clonal structure of the sample. We assume that CNAs appear in several rounds of clonal expansions. We assume that having many sub-clones is not likely, so we penalize each additional sub-clone with empirically chosen threshold (can be selected according to BIC criteria and number of regions under investigation). Then, we want to find an optimal set of clonal cell fractions from the pre-defined discrete set of 5%, 7.5%, …, 97.5%, 100% that will explain our CNAs in the best possible way, considering sub-clones as real only if they substantially improve the overall likelihood of the variants. For each cancer cell fraction *α* and each CNA we choose the best possible explanation (the state with the maximum score with the CCF fixed and equal to *α*). Then we investigate all potential combinations of sub-clonal fractions up to 5 clones. For each CNA we select one cancer cell fraction from the possible combination that explains this CNA in the best way.

When the likely sub-clonal structure is inferred, we run the calling procedure again, using only states that may arise from this restricted set of sub-clones.

We also developed a QC control for variants which is based on the fact that all the allele-imbalanced events are expected to have BAF signature different from the paired normal BAFs and CNA events in the regions of high variance or extremely low-covereage are, likely, false positives. The procedure is described in Supplementary.

## III. RESULTS

For evaluating analytical performance and comparison with existing methods, we analyzed the following data:

1. 473 WES sequenced tumor-normal pairs from CLL cohort (272 samples were enriched with Agilent 51MB kit and 169 samples with Agilent 71MB kit, only pre-calculated coverage counts and BAFs data were available for the whole cohort due to data migration issues). This dataset was already explored and described in [Puente et al., 2015] with the array-based methods, we re-analyse this cohort with WES and ClinCNV.
2. 251 TPS in-house sequenced tumor-normal pairs, sequenced with 3 different panels. Additionally, panels number 2 and 3 (in chronological order) had many samples sequenced with the low-input sample preparation protocol, which we separated from the others since their coverage profiles differ. Summing up, we had 5 different cohorts of TPS samples.

For comparisons using alternative tools we had:

1. 505 pairs from CLL cohort analysed with Affymetrix 6.0 SNP arrays (partially overlapping with our WES samples);
2. 80 raw alignment BAM files (40 tumor-normal pairs) from whole exome sequenced CLL cohort pairs (enriched with Agilent 71MB kit, initially selected for SNV calling algorithm study benchmarking
3. BAM files from in-house TPS samples

For array data analysis, we selected ASCAT ([Van Loo et al., 2010]) – a well-established tool for benchmarking of CNA calling pipelines that allows purity and ploidy estimation. For WES and TPS data analysis FACETS tool was applied due to its high performance and sophisticated algorithm that involves joint segmentation of two signals (from coverage at SNV positions and BAFs from heterozygous SNVs). CNV-Kit was used for comparison in WES data.

Newest available versions of tools were used: ClinCNV 1.16.0, FACETS 0.6.0, ASCAT 2.5.3, CNV-kit 0.9.6. Tools were evaluated using recommended parameters and pipelines. Internal quality control of variants was applied at the first step of the ClinCNV’s calling algorithm for TPS samples and for both steps for WES samples.

### i. CLL cohort results

The on- and off-target coverages were substantially different between two enrichment kits fig. 3.

**Figure 3:**
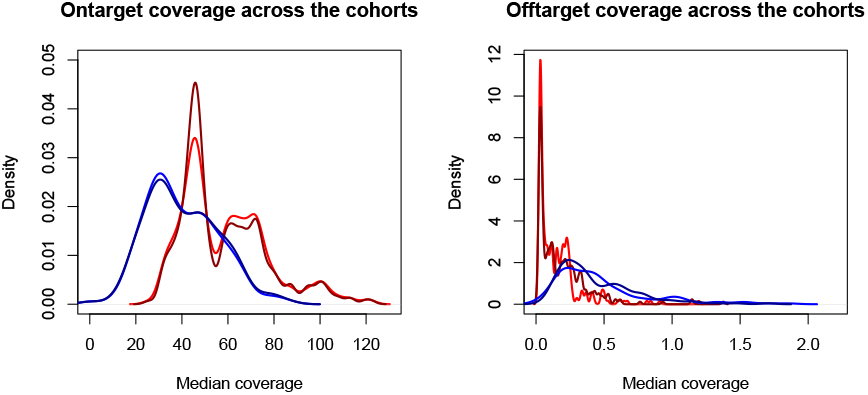
Median coverage plots. Dark red and red denote Agilent 51 tumor and normal, dark blue and blue – Agilent 71 tumor and normal. Average coverage is 15-20% higher than median coverage. As can be seen, on-target coverage for Agilent 51 was higher while for Agilent 71 off-target coverage becomes more valuable signal.

Six samples were excluded after the manual examination (described in supplementary) due to likely mislabeling or large stretches of zero covered genomic regions. The second part of QC control was done automatically. Samples with the coverage variance bigger than one were excluded automatically. We checked if a sample had a lot of false-positive events detected and either excluded such a sample from the analysis or removed likely false positive variants. In order to determine the level of false positives, we took all events where a major allele is not equal to a minor allele and found how many such CNAs (that contain at least one SNV) showed no significant deviation in BAF frequency (p-value > 0.5) since we expected deviation in BAF for imbalanced events. If the amount of likely false positive imbalanced variants was bigger than 10% of the overall number of detected CNAs, we said that the false discovery rate in this sample was too high and we discarded it. 20 samples (less than 5%) from our CLL cohort did not pass this check and were excluded. If the amount of likely false positive variants was bigger than 5%, we retained the sample, but removed all imbalanced events with FDR corrected p-value bigger than 0.05 to control FDR at least for copy-number imbalanced events (balanced events are relatively rare except for high ploidy samples). We did nothing if a sample was of good quality and less than 5% likely false positive results were found. We did not filter all imbalanced events with q-value bigger than 0.05 since for the short or low CCF true positive variants q-values may be bigger, but often these events were real. It is worth to mention that many false positive variants with increased/decreased coverage but an absence of any shift in BAFs were observed in other datasets sequenced with the high coverage and modern enrichment kits, so this phenomenon can not be explained by the unusual sequencing technical conditions of the investigated cohort.

21 out of 473 samples were marked as QC-failed and re-sequenced in the previous analysis of CLL data. They were included in our test cohort in order to check if these samples will be recognized by ClinCNV, and 11 of them were filtered out.

After these two steps, the QC-filtered cohort included tumor DNA sequenced from 433 samples. Only samples without QC issues were used for comparisons.

**Figure 4:**
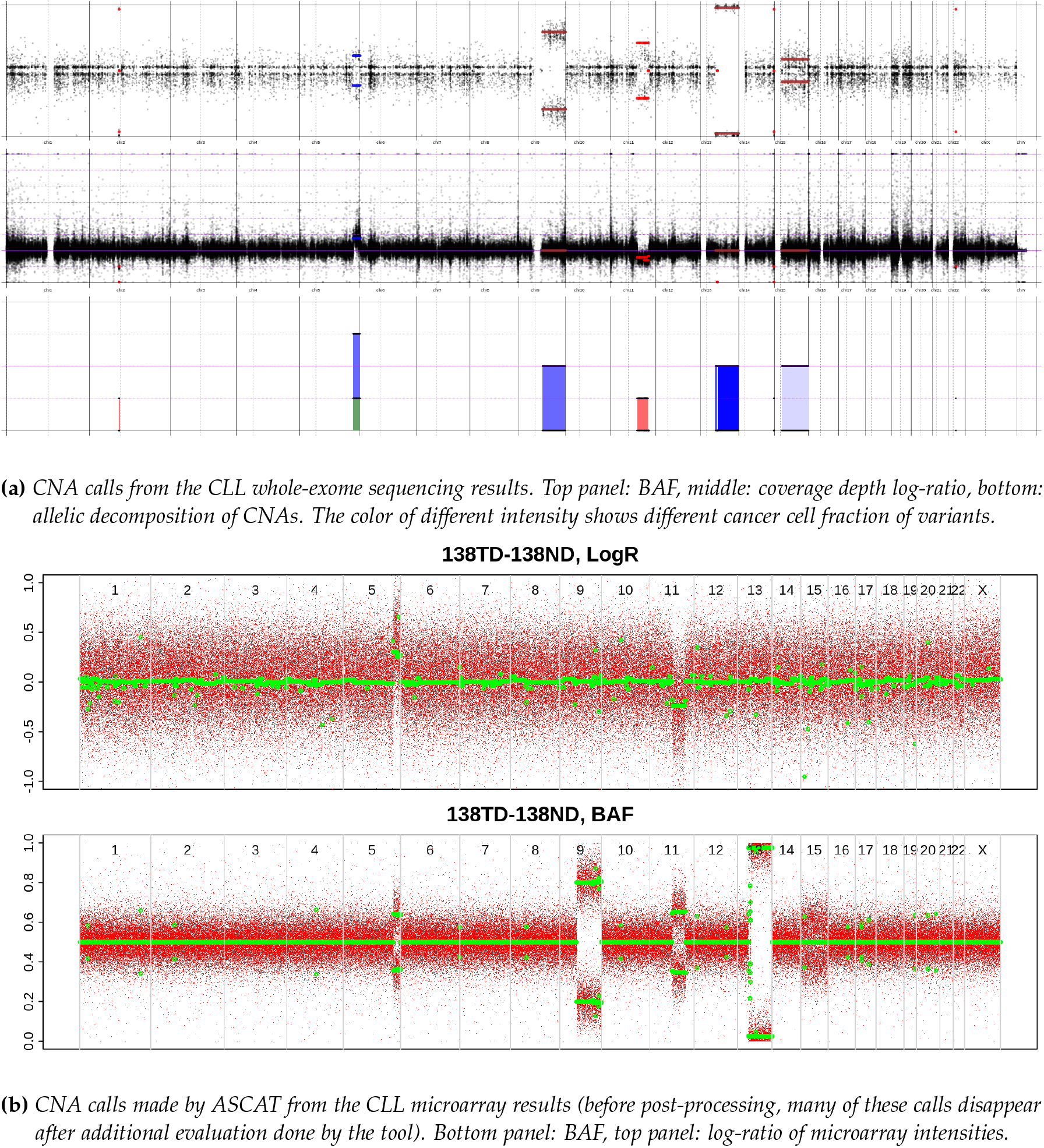
CNA calling results in the same CLL sample, obtained by ClinCNV in WES data and by ASCAT in microarray data. Due to the presence of several clear clones ASCAT mistakengly counted this samples as having ploidy 4. The LOH at chromosome 15 was not called by ASCAT, even if it can be recognised visually. 3 clones were identified by ClinCNV at cancer cell fractions of 97.5%, 60% and 17.5%.

#### i.1 Overall statistics of CLL callset

Size distributions of detected variants are shown in fig. 5.

**Figure 5:**
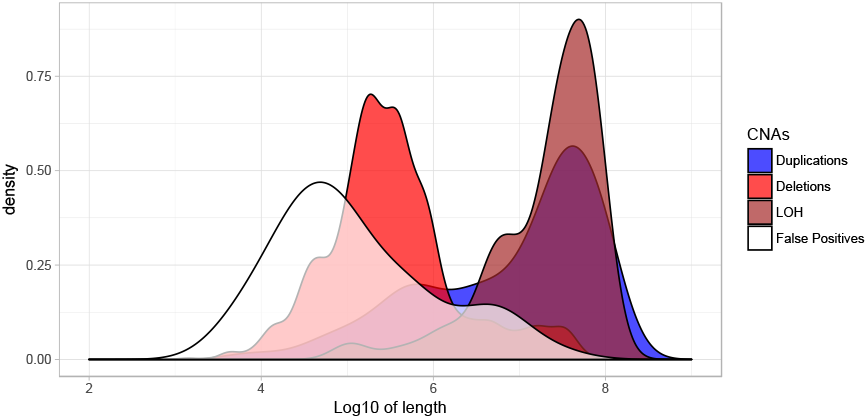
Length of detected variants, by type.

Distribution of cancer cell fraction of detected variants against length of variants is provided in fig. 6. As can be seen, the distributions are shifted to the top-right corner. It can be hypothesized that it is not only due to the abundance of variants with the large lengths and high cancer cell fraction, but also due to the power limitation of detection of short sub-clonal variants.

**Figure 6:**
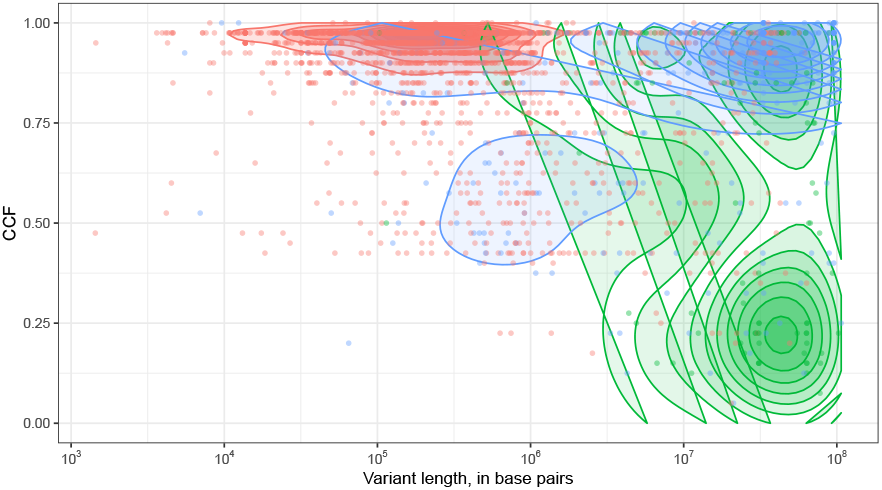
Distribution of cancer cell fraction of detected variants against length of variants. Blue dots denote duplications, green – LOH, red – deletions. Density contours are also shown.

#### i.2 False Discovery Rate

Since not all the calls from ClinCNV were present in ASCAT’s output, we came up with the procedure for autosomal CNA validation using array data pre-processed and normalized by ASCAT. The main idea was to create two distributions of p-values: a null distribution from simulated CNAs and a distribution of p-values from the real results. Having both, we could estimate FDR: in theory, two times the number of p-values, bigger than 0.5, should give us an estimate of FDR, but we decided to generate a null distribution in order to check if it is actually uniform (described in Supplementary).

P-values from real ClinCNV’s CNA calls were compared with the null distribution (fig. 7). For the analysis of LOH variants, we included BAF-based p-value only; for deletions and duplications, we merged p-values from array intensities and BAFs with Fisher method where possible. However, we have noticed that our null distribution was not strictly uniform. Hence, the doubled number of p-values which were bigger than the median of negative control p-values gave us a more accurate estimate of overall FDR.

**Figure 7:**
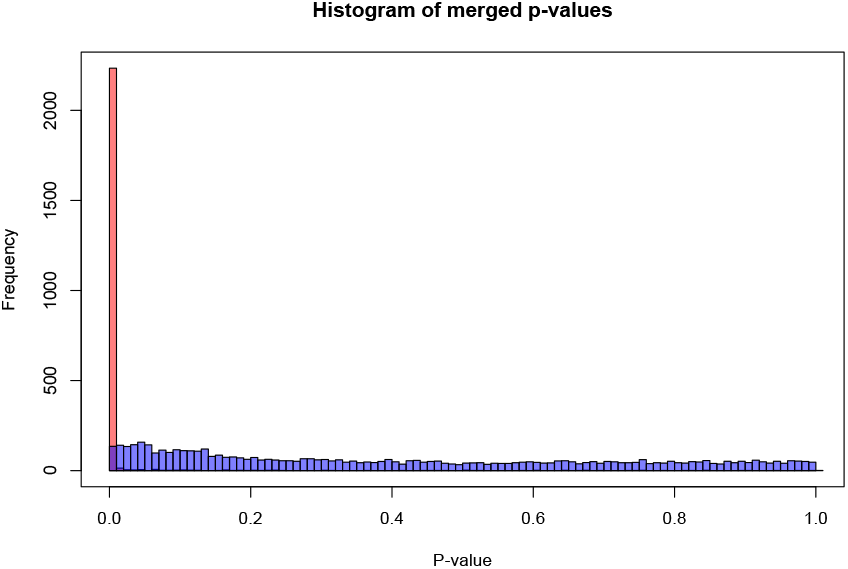
ClinCNV calls p-values (red) and simulated calls p-values (blue).

As an additional FDR check, we filled the contingency matrix of deletions and duplications, distinguishing between deletions and duplications that show positive/negative intensity in arrays. If ClinCNV called variants randomly, we could expect the array intensity to be independent of the CNA type called – array intensity may be shifted positively and negatively for false positive deletions or duplications with an equal probability.

Overall 2464 CNAs were detected in QC passed samples. 2341 CNAs were suitable for validation (contained at least one marker from arrays). Median of p-values from simulated CNAs was equal to 0.346, and 34 CNAs detected by ClinCNV had p-values bigger than this median. Thus we could conclude that false discovery rate of our callset was 2 · 34/2341 ~ 0.029. Analysis of the contingency matrix showed that 20 deletions and 13 duplications detected by ClinCNV showed discordant median array intensity, respectively. Thus we concluded that false discovery rate in copy-number imbalanced events was equal to 2 · (20 + 13)/(2250) ~ 0.0293.

#### i.3 Sensitivity comparing to arrays

We compared the ASCAT callset from microarray data with the results of ClinCNV in WES data in a similar manner. Among 3787 variants detected by arrays and containing a sufficient amount of markers in WES, we found out that we potentially missed 490 CNAs. Analysis of events with copy-number changes (deletions and duplications) detected by ASCAT but not found by ClinCNV showed that 1974 of them showed concordant direction (variant detected as deletion showed lower coverage in WES and higher for duplications) and 1425 showed discordant direction, which gave us estimation of 549 potentially missed CNAs (which is bigger than estimation of 490 due to low power of detection in some regions in WES comparing to arrays). However such evaluation also showed us that false discovery rate of ASCAT in array data was high – ClinCNV detected 1708 CNAs matching to 1321 array results and ASCAT detected 3787 unique variants in arrays. Only around 550 of them were also observed in NGS data, so we can roughly estimate FDR of ASCAT as bigger than 50%. We hypothesize that ClinCNV may detect such variants using more relaxed thresholds, which will lead to a higher false discovery rate. Such calculations led to the estimation of the sensitivity of ClinCNV (using variants that span at least 5 WES markers) of approximately 1 − 550/(550 + 1319) ~ 0.7 (comparing to array-based method).

The purities of samples estimated by ClinCNV (as maximum cancer cell fraction of variants) and ASCAT were compared in fig. 8. Aberrant cell fraction estimated by ClinCNV for some samples was close to 1 while much smaller as estimated by ASCAT – after manual checking of these samples we concluded that such divergence in estimations mostly happened in samples where arrays had not found any large significant CNA while for almost all the samples aberrations in immunoglobulin genes regions occured and were detected by ClinCNV.

**Figure 8:**
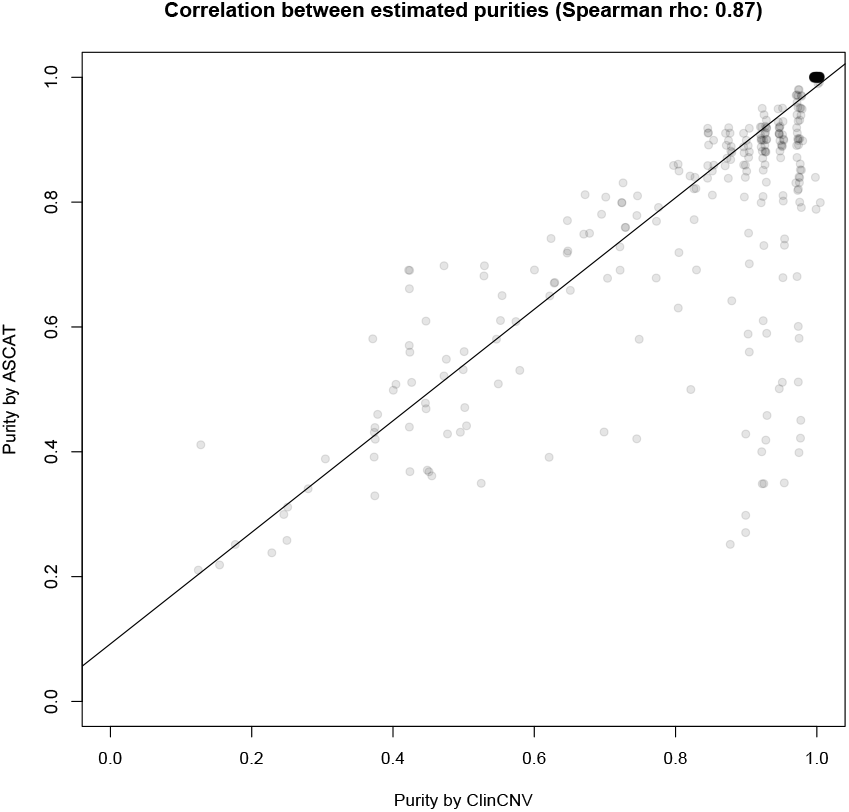
Purities (ClinCNV’s estimation vs. ASCAT estimation). Spearman correlation: 0.874.

#### i.4 Comparison with other tools

We have validated FACETS calls from 40 WES sample pairs using the same procedure as ClinCNV. No QC failed samples were present in this dataset, and one variant was removed at the postprocessing step from the ClinCNV’s callset according to the QC guidelines provided above. FACETS calling was performed using ten reads as a minimum depth, and cvalue quality threshold was chosen equal to 150, according to the authors’ recommendations.

Out of 190 autosomal raw calls from FACETS 152 had at least one array marker within the borders and thus were suitable for validation. Seventeen variants had p-values bigger than the median of a negative control, so using our methodology at least 2 · 17 = 34 variants were false positives. Also, 19 deletions showed positive array intensity shift; four amplifications showed a negative shift, and 93 and 17 deletions/duplications, respectively, showed the correct direction. Thus, FACETS FDR was estimated as 0.22 for all variants from p-values, 0.35 for variants with change in copy-number from the contingency table.

CNV-kit evaluation was complicated since it does not detect LOH events by default, and it does not estimate purity by itself. However, our data were obtained with FACS-sorted blood cells sequencing results, so we may assume that the purities of the samples were close to 100%. CNV-kit detected 137 autosomal copy-number imbalanced events from these 40 samples. 132 variants were suitable for validation with arrays, and the estimated FDR was equal to 0.56 based on p-values and 0.77 based on the contingency matrix. The potential sources of differences in FDR estimated with two methods are described in Supplementary.

Using the same 40 tumor-normal pairs, we were able to validate 179 autosomal variants from 181 ClinCNV raw calls. None of the detected deletions/duplications had an incorrect direction of array intensity shift. One variant had p-value bigger than the median of the null distribution, thus, ClinCNV’s FDR in this sub-group may be estimated as 2 · 1/176 = 0.011.

As for concordance between methods, we chose CNAs that had p-values less than 0.01 in array BAFs or intensities as True Positives and considered two variants as matching if they overlap for at least 50% of the length of the smallest variant. The number of True Positives in ClinCNV was equal to 178, in CNVKit 52, in FACETS 109 (table 1). 122 True Positive variants from ClinCNV were mapped into 97 True Positive variants from FACETS. Such difference was not caused by over-segmentation, but rather, it was due to the fact that ClinCNV analyses each chromosome arm separately so aneuploidies (such as chr12 duplication, frequent in CLL tumors) were represented as two variants in ClinCNV’s output. 50 ClinCNV True Positive variants matched with 39 variants from CNVKit. ClinCNV detected 56 (52 if we consider closely located variants as one) real CNAs not present in FACETS calls while 12 variants from FACETS were not present in ClinCNV’s callset. CNV-Kit detected 13 variants not present in ClinCNV callset while 128 CNAs from ClinCNV’s callset were not detected by CNV-Kit (9 of which were LOH events nondetectable by CNV-Kit using the default pipeline). Concordance between FACETS and CNV-kit was even lower: 32 variants from FACETS were matched with 30 variants from CNV-kit.

**Table 1:**
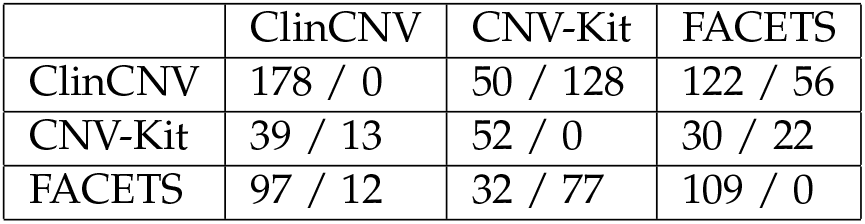
Table of concordance. First number denotes number of True Positive CNAs from the callset matched with another True Positive callset, second number denotes number of TP events detected by the tool uniquely comparing to another callset.

### ii. Targeted panel sequenced cohort results

For our TP-sequenced cohort, we had no array data to validate so we could only compare results from different tools. CNV-kit was excluded from the comparison due to the necessity of third-party tools for purity estimation. We have selected only samples with tumor content determined by a pathologist as bigger than 20% since TP-sequenced samples of lower purity are usually discarded by molecular tumor boards and thus accuracy of calling in such samples is not crucial. We additionally excluded samples where pathology data on purity was not available and ended up with 216 tumor-normal pairs from 251 initially sequenced samples.

We say that a variant from FACETS matched a variant from ClinCNV if their overlap was bigger than 50% of the smallest variant length and vice versa. 3654 variants from FACETS were matched with 5772 variants from ClinCNV. Again, such a difference was likely due to the fact that ClinCNV operates with chromosome arms while FACETS operates with whole chromosomes and ClinCNV utilizes off-target reads while FACETS is mainly concentrated on working with the targeted regions. FACETS detected 564 variants not matched with ClinCNV variants while ClinCNV detected 1196 variants not presented in FACETS callset. On average ClinCNV detected 2.9 variants more than FACETS per sample (median: 1) which may be a consequence of higher sensitivity in off-target regions.

Comparing purities estimated by these two methods (estimated as maximum cancer cell fraction of detected CNA for ClinCNV, 0% if no CNAs were detected, and extracted as a designated value from FACETS calls), we found that despite the good concordance between methods in general, a large number of samples have higher purity in ClinCNV estimations (bottom right corner at fig. 9).

**Figure 9:**
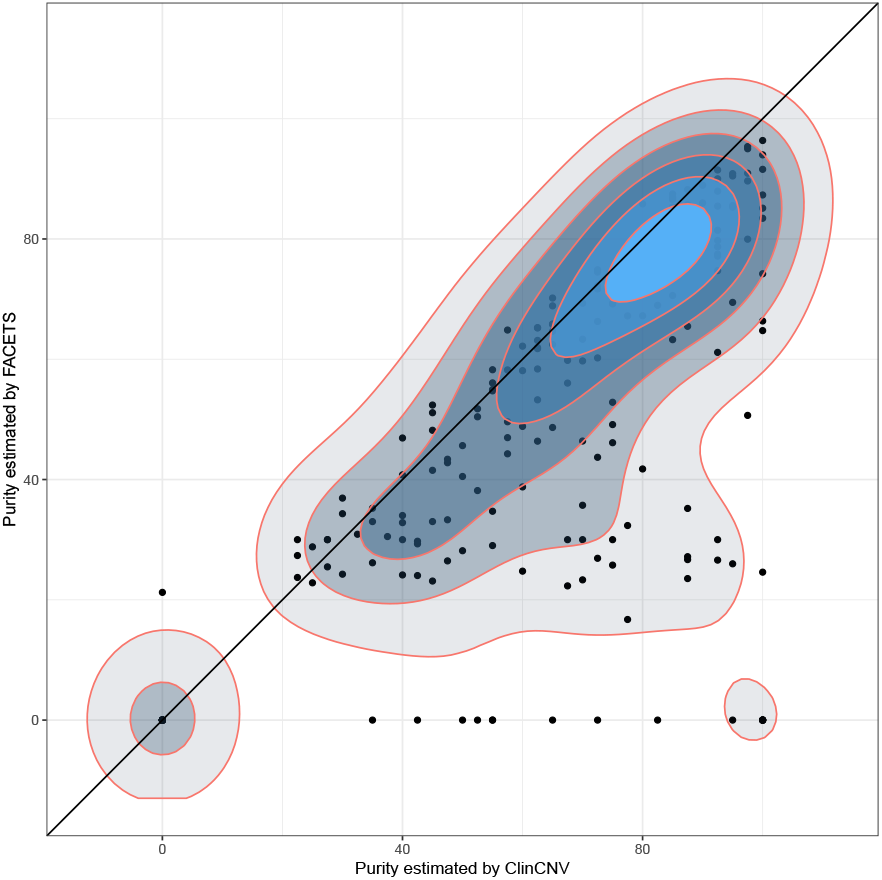
Purities of targeted panel sequenced samples estimated by ClinCNV and FACETS.

We hypothesized that it might occur due to the fact that ClinCNV penalizes higher copy numbers as less realistic while FACETS finds an optimal solution without taking this into consideration. In order to check which method is better, we decided to compare purity values with pathology estimation even if pathology estimation is regularly reported as being inaccurate. We have found that even if for some samples estimations from FACETS were more accurate, for a large amount of samples FACETS underestimated purity, potentially, due to its optimization process that does not take the lower prior probability of large copy-number changes into account.

To make it more obvious, we concentrate on samples with more than 25% discrepant purity between ClinCNV and FACETS and show their purities in comparison with pathologists estimation (fig. 10).

**Figure 10:**
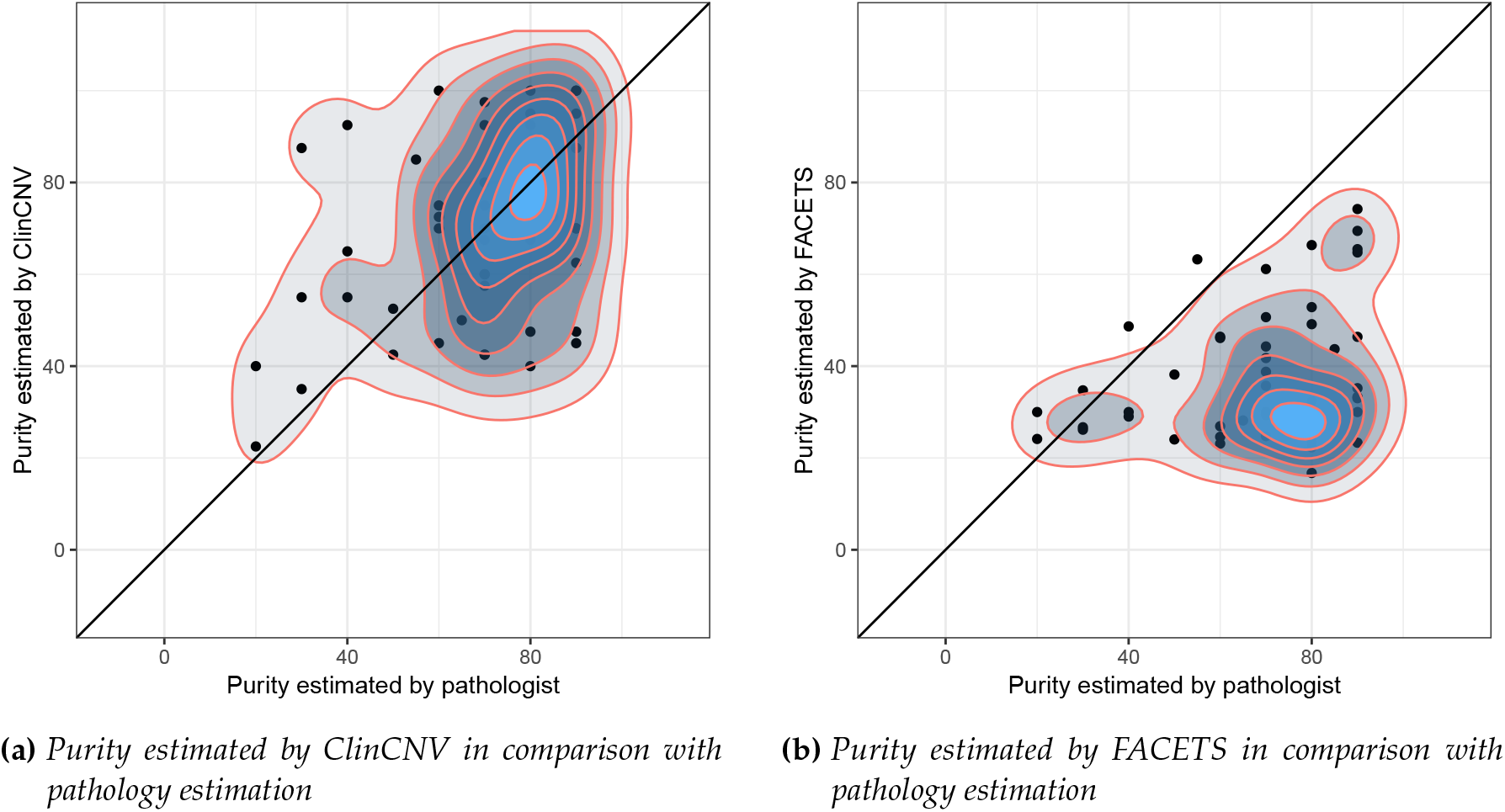
Purity of discordant samples (more than 25% difference in purity estimations) of ClinCNV and FACETS in comparison with pathologist estimations.

Thus, the median difference between ClinCNV predicted value and pathologist tumor content estimation was −5% (mean −7.5%) and for FACETS median difference was −19% (mean −23%). Using MAD as a measure of variability, we can estimate that ClinCNV median absolute deviation was 15% while for FACETS it was 25% (likely, due to bi-modality of the distribution). Otherwise, methods showed a similar degree of variability comparing to pathology estimation.

## IV DISCUSSION

We have developed a new powerful approach for CNA detection in paired tumor-normal NGS data and showed its superior performance compared to other tools for CNA detection in large cohort of CLL samples. We also showed good concordance with existing method in cohort of cancer-gene panel targeted sequenced samples. We performed callset analysis using Cancer Genome Interpreter and showed that CNAs were often annotated as biomarkers of response, resistance or toxicity as a response to particular drugs in many samples (Supplementary Materials). Thus, CNAs calling using ClinCNV and annotation of the calls can be routinely done and reported together with other biomarkers such as point mutations or indels, tumor mutational burden, gene expression of specific markers, etc.. ClinCNV may provide an extensive summary of CNAs in tumor genomes which is of extreme importance for the healthcare.

## Supporting information

Supplementary Methods and Results

