## Supplementary Methods and Results for "ClinCNV: novel method for allele-specific somatic copy-number alterations detection"

### I. Supplementary Methods

#### i. Sample level QC control

As a simple sample level quality control, we always filter out samples for which robustly estimated standard deviation of log-ratios in autosomes is greater than one because such a huge variance always indicated technically failed samples and not a complex CNAs pattern in the test data. ClinCNV allows analysis of such samples, but only if their IDs are specified explicitly as an input parameter which means that the investigator is aware of potential QC issues.

##### i.1 Selection of SNVs for BAF analysis and CNA detection.

In order to facilitate in-depth analysis of allelic imbalances, we use bi-allelic single nucleotide variants, namely their BAFs (ratio between the number of reads supporting the alternative allele of an SNV to the overall coverage of the position). We use only well-covered SNVs ($>20$ reads in tumor and $>60$ reads for both tumor and normal) since low-covered SNVs tend to be erroneous and provide a small amount of evidence of CNA while increasing computational time. We use only positions that are likely heterozygous in the normal tissue (BAF between 0.4 and 0.6). Additionally, to determine whether the particular SNV is heterozygous or not we take into account "inherent biases due to differential mapping affinity between the reference and the variant allele" ([Shen et al., 2016]). To correct this bias, we calculate the median of the BAFs. Then we only include SNVs in the analysis in which allele-specific coverage ratio was not rejected by the two-tailed Binomial test at the 0.01 level of significance, using the previously determined median as the expected probability. Finally, we remove all SNVs that form clusters (located closer than 20 bp from each other) in the genome since such SNVs are enriched for false positives and not independent (present in the same read).

#### ii. Purpose of the in-direct estimation of variances

The indirect way of variance inference described in the paper may help in 3 ways:

*Corresponding author

1. It is not affected by recurrent Tumor CNAs which can be crucial for the analysis of cancers with known recurrent CNAs (such as short deletions in chr13q14 in chronic lymphocytic leukemia malignancies) since we used only the data from healthy tissues for individual regions' variances estimation;

2. For off-target normalized counts we often see that directly observed variability of a particular region in log-ratio data is much smaller than the value predicted by the method above. It means that these off-target regions are highly affected by non-diploid polymorphisms, and that diploidy assumptions may be violated. These regions are subsequently excluded from the analysis if the observed and predicted variance ratios differ by more than three standard deviations.

3. An estimated variance in general is usually lower than the variance observed directly from the data, especially for off-target reads. It is caused by the large variability in the proportion of off-target reads for different samples. The number of on-target reads is relatively stable across a particular cohort of samples.

### iii. Detailed description of the segmentation and calling algorithm

Having data normalized and parameters estimated we start calling procedure (fig. 1). At first, we define a set of potential copy-number changes that may occur in a tumor sample at each allele. In the beginning, we investigate major and minor alleles under assumptions of:

1. Discrete set of the potential clonal fraction (starting from 5 to 100% with the step of 2.5%, further increase of resolution did not improve calling for both $30 - 60$x WES and $> 200$x TPS samples);

2. Minor copy number from 0 to 4;

3. Major copy number from 0 to 30: major and minor copies together are limited to 30 copies (we trim our data so all the values that show higher copy number change will be at 30 copies with $\sim$100% CCF model).

Having probabilistic models and set of states $S$ we can calculate likelihoods of each data point $x_1, \ldots, x_n$ under all these models: $\mathcal{L}(S_a|x_i) = p_{S_a}(x_i) = P_{S_a}(x_i)$. We fill a matrix of size $|S| \times |G|$, where $|S|$ denotes the number of states and $|G|$ is the length of the genome with the corresponding likelihoods. Ordering of genomic regions must be preserved in such matrix, in other words, the first column of the matrix of likelihoods should contain likelihoods of data points obtained from the most "left" (upstream) part of the genome or its target of interest, the second column should contain the likelihood of the data point from the next genomic window and so on. It is usually possible to define a baseline copy-number state: it is a diploid state for human or mice autosomes, or a haploid state is a baseline for males' sex chromosomes. We denote the probabilistic model corresponding to such a baseline state as $S_b$. The problem of finding one CNV in such a matrix may be formulated as:

**Problem 1** *Having matrix of likelihoods of datapoints under different states and baseilne state $S_b$, identify a pair of indices $i, j$ and state $S_a \neq S_b$ such as $1 \leq i \leq j \leq |G|$ and*

$$\arg\max_{i,j,S_a} \frac{P(S_a|x_i, \ldots, x_j)}{P(S_b|x_i, \ldots, x_j)}$$

Informally, we need to identify a probabilistic state, and a start and end position of a genomic segment that shows the highest evidence that a model different from the baseline fits the observed coverage depth signature better. Or in brief, we are looking for a segment with the highest evidence of being in an alternative state. Naive computational solutions to this problem that require checking of all possible $i, j, S_a$ are not computationally feasible since it takes $\mathcal{O}(|S||G|^2)$ operations. Even assuming independence of random variables, we would need to calculate $\frac{P(S_a|x_k) \cdot P(S_a|x_{k+1}) \cdot \ldots \cdot P(S_a|x_l)}{P(S_b|x_k) \cdot P(S_b|x_{k+1}) \cdot \ldots \cdot P(S_b|x_l)}$ for

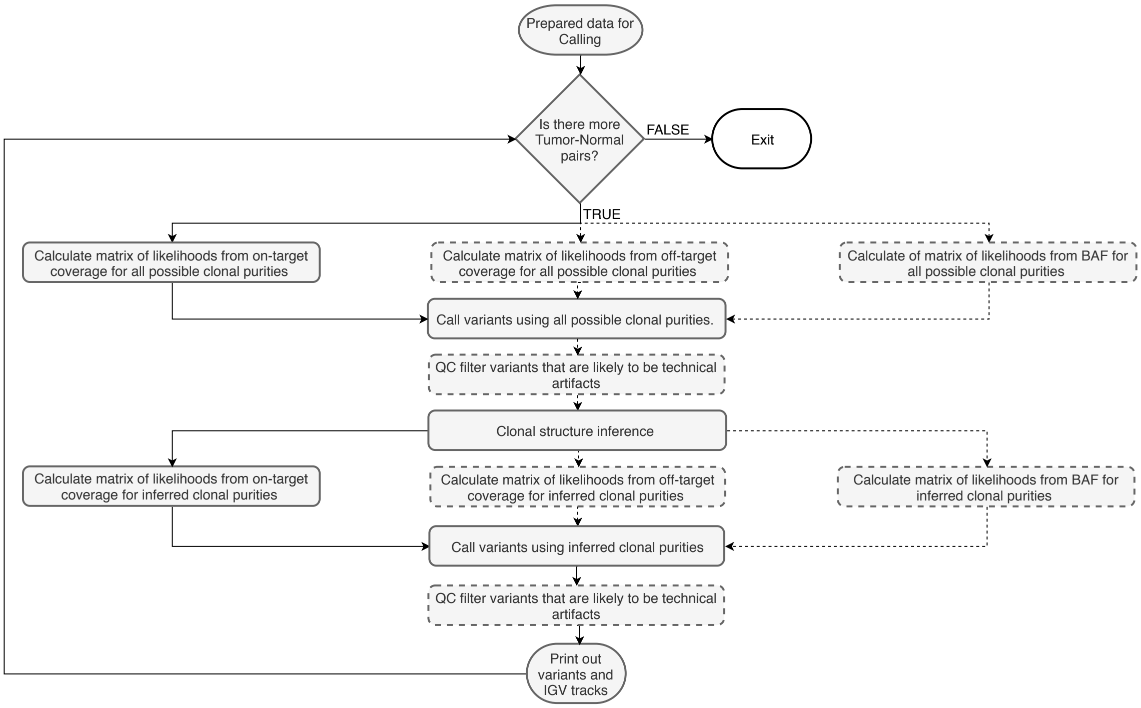


**Figure 1:** *Flowchart of CNA calling.*

each pair of $k, l$ such that $1 \leq k \leq l \leq| G |$ and choose $k, l$ such as value of this expression will be maximized.

To simplify the problem, we take logarithms of likelihoods so we will be able to sum them instead of multiplying (the example of log-transformed matrix of likelihoods is depicted on the top of fig. 2), so now we have a matrix of $l(S_a|x_i) = log(\mathcal{L}(S_a|x_i)$. Then, for each genomic region $i$ we can switch to likelihood ratio $\log \frac{\mathcal{L}(S_a|x_i)}{\mathcal{L}(S_b|x_i)} = l(S_a|x_i) - l(S_b|x_i)$ (in the middle of fig. 2, after "Log-likelihood differences of the models" header). This value is already meaningful – it shows how strong is the evidence of $S_a$ against $S_b$ if the log-likelihood ratio is positive and how strong is the evidence of $S_b$ over $S_a$ in case of negative log-likelihood ratio. If we want to find a segment and a state that shows the biggest evidence against $S_b$ we need to find a segment with the largest log-likelihood sum for each state and then choose the state where we found a segment with the largest value of log-likelihood sums. To find segments with the largest sums for one particular state, we can use a well-known maximum subarray sum algorithm that solves the following problem:

**Problem 2** *Giving a one-dimensional array of numbers $A, A[i] \in \mathbb{R}$, find indices $i$ and $j$, $1 \leq i \leq j \leq n$, such as $\sum_{k=i}^{j} A[k]$ is a large as possible.*

Kadane's algorithm [Bentley et al., 1984] solves this problem in linear time. We can apply Kadane's algorithm $|S| - 1$ times for each state except the baseline one and choose the segment and the state with the largest sum as an answer to the problem (on the bottom of fig. 2).

We reduce the number of steps to $\mathcal{O}(|S||G|)$ for solving **Problem 1** using the described procedure.

Whenever we have different signals (such as read depth and B-allele frequency), we can simply sum up matrices of likelihoods from different signal types and perform the same procedure, so it does not affect computational time except for the additional calculations of likelihoods. Models of any complexity may be applied for finding likelihoods without an increase of computational time on segmentation.

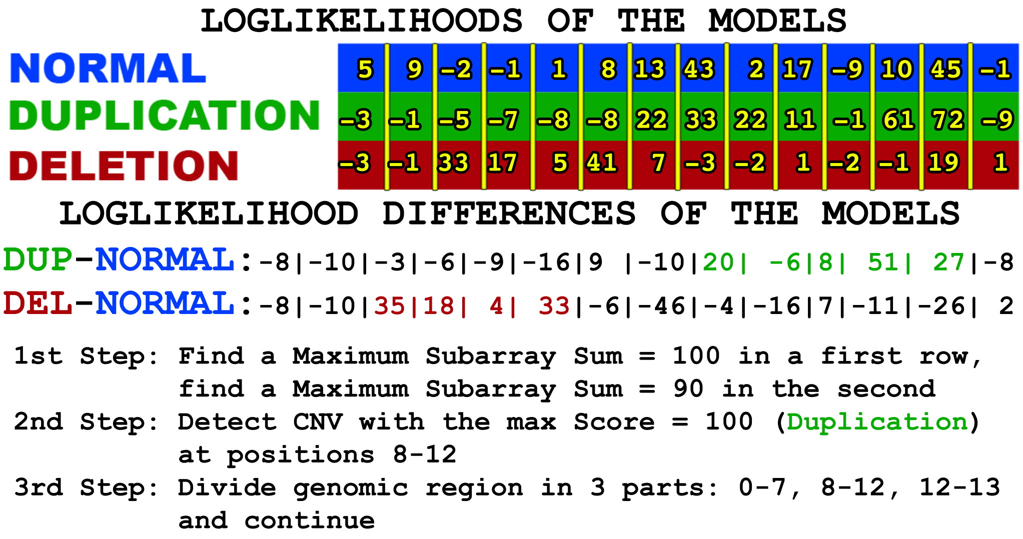


**Figure 2:** *Toy example on finding one CNV with the help of matrix of likelihoods.*

Such a procedure does not require a calling step afterward – everything is inferred by determining which $S_a$ is the best for the explanation of the observed data comparing to the baseline $S_b$.

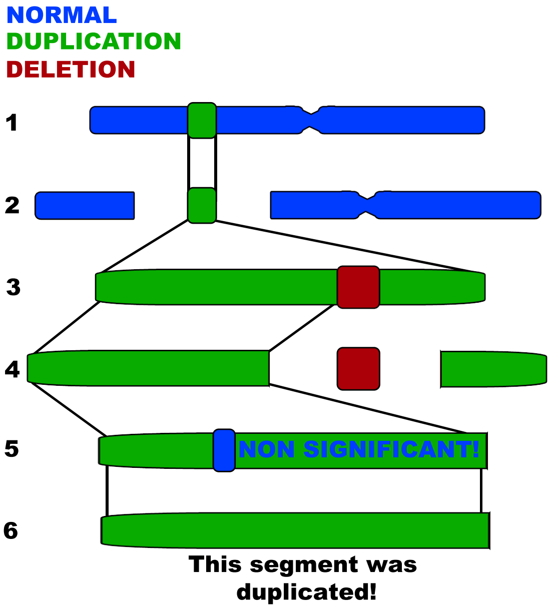


**Figure 3:** *One branch of the recoursive tree of the segmentation algorithm.*

Segmentation of a genomic piece into candidate CNV regions is done analogously to CBS (fig. 3). We find one piece that shows the presence of an alternative model at one step of our algorithm, and then we divide the initial genomic piece into three segments: one to the left, one to the right from the discovered segment and the discovered segment itself. We stop segmenting once the next detected segment fails to reach a significance threshold. If the found segment is significant but shorter than the predefined length, we decrease the evidence of an alternative state that comes from this region, and then we re-try to find a significant segment of sufficient length again. Once all the branches of the recursive tree report absence of significantly different genomic sub-segments, the algorithm collects results from all the endpoints of the recursive tree and form a resulting callset.

After the investigation of several datasets, we have found a CNA signature in BAF and read depth that was not correctly recognized by the described models. The coverage was slightly decreased in such regions, which may indicate a deletion in a small sub-clone, while BAF was largely shifted from the expected values. The most likely explanation to such pattern is that a deletion was followed by a duplication, but not in all of the tumor cells affected by the deletion. Alternatively, a LOH event occurred, followed by a deletion in less than 100% of cells that experienced the LOH event. In order to detect such events without over-segmentation of the calls, we had to introduce a “second round of CNA calling” even if ClinCNV does not reconstruct a timeline of somatic alterations. We added limited support of second-round CNAs with the following restrictions: 1) for interpretability, second-round CNAs may happen only in less than 50% of cells that experienced the first copy-number alteration, so in most cases we can neglect such changes; 2) second-round CNA can be only a simple deletion or duplication of one copy; 3) second-round CNAs are considered only after heterozygous deletions or simple losses of heterozygosity, since otherwise, the calling will become ambiguous. We add these states only after clonal structure inference due to significant increase in the number of states and thus the computational time of calling.

To correct for potential ploidy change we not only select BAF-balanced chromosomes' arms for normalization, but also try to separate tetraploid regions from diploid, using the assumption that the smallest possible BAF-balanced chromosome' arm copy-number is 2 (since, as mentioned in [Shen et al., 2016], long stretches of homozygous deletions most probably lead to cell death and thus are highly unlikely). In order to do this, we sub-select all chromosome arms with small deviations in BAF and calculate coverage baselines (medians of log-ratios of coverages) for all such arms. Then we iteratively merge all arms with differences in medians of their normalized coverage less than 2.5% and choose the smallest merged value that includes at least 10% of markers from BAF-balanced chromosome arms as the baseline.

Having all the parameters estimated and expected values for BAF and coverage ratio inferred, we perform calling, using the algorithm and states described above. We do not include unrealistic states (e.g., if all the coverage values are small, then no high copy-number event of high purity can be expected). Usually, we have more than 2000 states used for the initial calling.

After we have finished the first round of calling using all possible purities, we calculate the likely sub-clonal structure of the sample. We assume that CNAs appear in several rounds of clonal expansions (otherwise, calling may be stopped at the first step, however it may complicate the interpretation of sub-clonal CNAs). We assume that having many sub-clones is not likely, so we penalize each additional sub-clone. Then, we want to find an optimal set of clonal cell fractions from the pre-defined discrete set of 5%, 7.5%, . . . , 97.5%, 100% that will explain our CNAs in the best possible way, considering sub-clones as real only if they substantially improve the overall likelihood of the variants.

We do it in the following way: for each cancer cell fraction $\alpha$ and each CNA we choose the best possible explanation (the state with the maximum score with the CCF fixed and equal to $\alpha$). Then we investigate all potential combinations of sub-clonal fractions up to 5 clones. For each CNA we select one cancer cell fraction from the possible combination that explains this CNA in the best way. Each additional clone is penalized by empirically chosen value (this input parameter may vary from 100 to 500, the bigger penalty suggests a smaller number of sub-clones detected by ClinCNV). Such penalty may be set equal to BIC penalty for additional parameter depending on the logarithm of the overall number of genomic regions, but in reality, better results are achieved with more strict penalization. Normally we set the additional clone penalty to 300, but it may be too strict for, e.g., panel sequencing data from a small panel of tens of genes, and thus has to be decreased. ClinCNV does not fully investigate the landscape of complex CNAs that have nested structure (e.g., duplication in $X$ percents of tumor cells followed by a deletion in $Y < X$ percents of tumor cells within the borders of the duplication) so such events can be detected as separate clones, and additional analysis has to be performed in order to investigate evolutionary history of such CNAs and "real" sub-clonal structure.

We output CNAs together with their scores and diagnostic information, such as 95% confidence intervals of on- and off-target coverage in tumor sample, median of tumor BAFs that are above and below the expected BAF from the normal sample which is helpful for checking of complex variants (such as overlapping CNAs from different clones, as described above), q-value of BAFs within the variant and overall q-value which we obtain by merging p-values from two sources (coverage-based signal and from BAFs) with Fisher's method, followed by BH FDR correction. The CCF of the biggest detected sub-clone is reported as the purity of a tumor sample. False positive results may appear, especially in low-purity samples, and this information may help to catch such variants during the post-processing.

#### iii.1 QC control

Complex quality control procedure was proposed. It takes into consideration potential markers of false-positive variants such as extensive variability in coverage or absence of evidence from SNVs' BAFs or presence of zero covered regions. Such QC control may be turned off for achieving maximum sensitivity, performed only at the first step of the algorithm (inference of clonal structure) or at both steps of the algorithm for the maximum specificity of calls. We were able to achieve the best results applying QC control at both steps for exome samples and applying this filter only for clonal structure inference for panel sequencing samples since the SNV signal is much more sparse in the latter type of data. QC control works in this way: we filter all variants for which the within-segment variability is greater than three times the overall sample's variability. We also filter out "homozygous deletions" that show the presence of normal coverage within the borders of a variant, so the median coverage of the segment is 0, but some regions have coverage log-ratio similar to the expected for the diploid regions. During the test runs of our tool, we noticed that copy-number alterations of small purity (less than 40%) in 30-60x covered WES samples (increased/decreased stretches of normalized coverage ratios) are often false positives. The coverage change is present and highly significant, but neither BAF signal changes nor evidence of such events was found in array data. For this reason we introduced a quality control procedure for each CNA call if the cell fraction affected by such a call is smaller than 40%. We create a baseline set of deviations in BAFs based on chromosome arms used for initial normalization since pre-selected chromosomes have the smallest proportion of deviations in BAF comparing to the normal tissue. For each CNA that is subject to additional filtering, we perform the Wilcoxon test using BAF deviations and discard CNA if its p-value is bigger than $10^{-4}$ (which means – the null hypothesis that BAFs of this CNA is equal to the expected diploid BAFs was not rejected). Variants with small purity that harbor less than 5 SNVs are discarded since no information is available for the quality evaluation. Finally, we retain the filtered variants if they meet previously described BAF deviation criteria (the BAF pattern is different from the expected). For example, the high within-variant variance of coverage which may occur due to technical reasons, but it also may be caused by undersegmentation of 2 relatively short closely located variants, thus, it is better to keep this CNA in a callset if it shows large deviation in B-allele frequency. BAF signal usually remains unaffected by coverage artifacts so, if BAF indicates a presence of a variant, then this variant is likely real even if its coverage is noisy.

### II. Supplementary Results

#### i. Another type of plot generated by ClinCNV

We provide several visualization options as, e.g., fig. 4, which may be easier for some professionals to interpret rather than plots with coverages and BAFs.

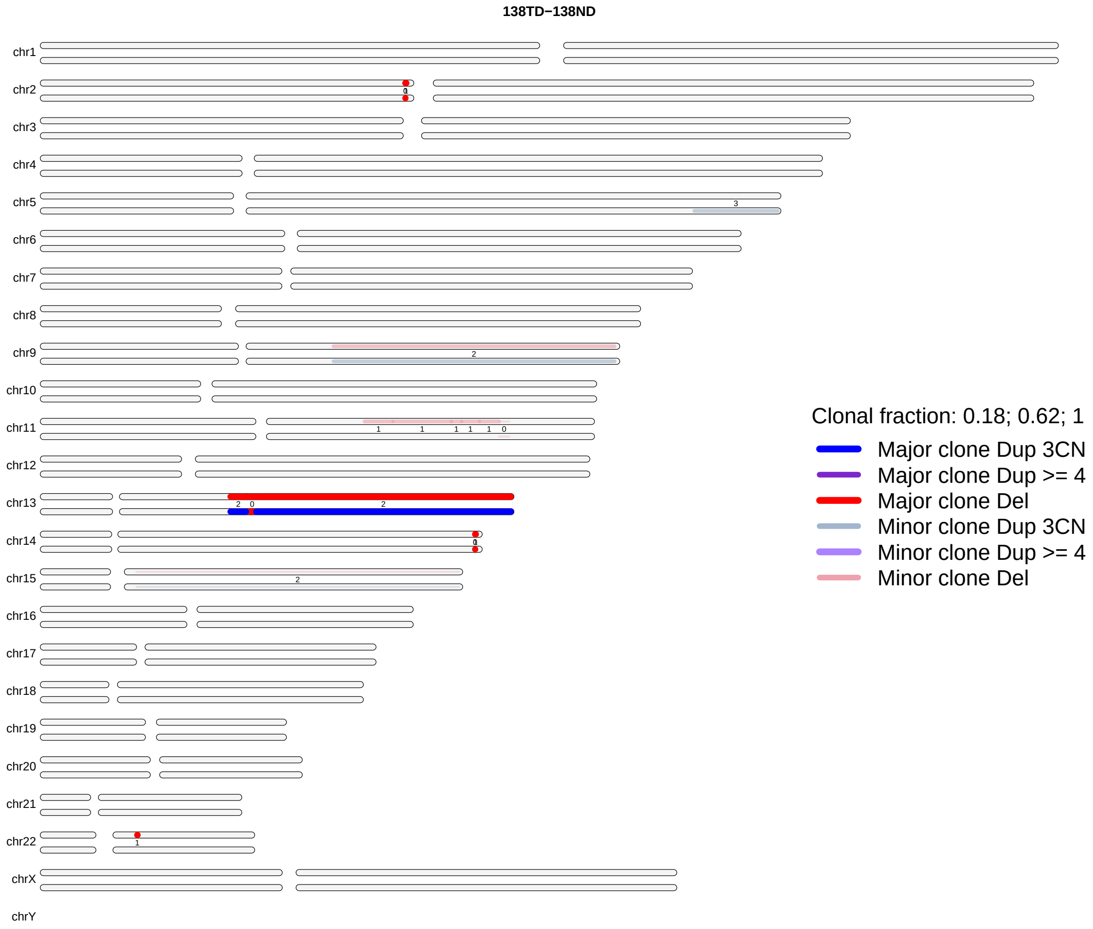


**Figure 4:** *ClinCNV provides several visualization options, depending on the intended purposes of the demonstration.*

### ii. Differences between research-purpose CNA calling and CNA calling as a clinical routine

CNA detection in tumor-normal pairs can be performed for research purposes (e.g., for finding genomic alterations responsible for targeted treatment outcome in clinical trials) as well as in routine clinical practice (common goal of the sequencing of tumor samples in clinics is to find alterations responsible for drug sensitivity, resistance or increased toxicity). The main differences between research and clinical approaches are:

1. Sizes of cohorts sequenced with the same sequencing panels and thus suitable for joint analysis. Large consortia such as TCGA generated cohorts of thousands of uniformly sequenced cancer samples while in clinics different custom panels are frequently used and the amount of samples uniformly sequenced with the same panel is usually of the order of tens or hundreds of samples.
2. In clinics, each calling result may be (and should be) manually examined and assessed by an experienced data analyst while it is not suitable for research samples.
3. Typically it is acceptable to filter out a

significant percentage of low sequencing quality or low purity or extremely complex tumor samples in research, but in clinics, each sequencing result often has to be analyzed since it is often not possible to re-sequence a low-quality sample due to increasing financial costs, limited time, complexity of biopsy extraction. Certainly, samples of extremely bad quality have to be discarded even in clinical practice, but filtering particular low-quality CNAs instead of filtering the whole samples is preferred in clinics. However, a specialist usually does not have infinite time for the resolving of one case, and the output of the tool should allow the analyst to prepare a CNA report in less than one hour.

Thus, quality control procedures should be automatized for research purposes, and summary statistics for manual assessment of each copy-number alteration have to be provided for clinical practice. ClinCNV does both, less strict filtering is usually applied for the clinical samples, however, no manual post-processing was done for the samples reported in this study.

### iii. CLL cohort results

**Sample-level Quality Control**

A sample quality assessment was done using the following procedure. We do not automatically check if samples were mixed (simple mislabeling or wrong pairing of normal and tumor data from different patients or a normal sample contaminated with a tumor tissue). Wrong pairing can be seen from a large number of more or less uniformly distributed short LOH regions and usually normal sample contamination can be seen in the BAF track of a normal sample. We were able to identify 4 samples (IDs: 1277, 1364, 1297, 1365) that demonstrated strong evidence of being mixed – almost whole genome was determined as LOH in such samples with many sparse heterozygous positions in-between. 2 out of such 4 samples were seemingly normal in array-based results, but the sex chromosomes did not match data from WES, which shows that it is more likely to be mislabeling than an unrecognized complex change (e.g., high ploidy). Two samples out of these 4 showed the same pattern in arrays. Another common pattern of low sequencing quality was many comparatively short regions of zero coverage. ClinCNV automatically filters them out if there are more than five homozygous deletions of length less than ten regions, but sometimes such 0-coverage regions are longer, and such samples have to be examined manually. Due to this consideration we excluded two samples that showed large 0-coverage stretches. These 0-coverage regions were false positives (presence of genomic material there was confirmed by microarrays).

#### iii.1 Detailed description of the FDR estimation procedure

**Checking WES-based results in Array Data: Null distribution**

For the null distribution, 10 CNAs per sample – five deletions and five duplications – of random size, ranging from 1 to 100kb, were introduced into ClinCNV's callset at positions not affected by ClinCNV's CNAs. P-values from permutation tests were calculated for BAF and intensity signals if at least one array marker was inside the borders of a CNA. For each CNA with $n$ markers inside, we sampled $n$ markers from CNA-free regions 1000 times and calculated p-value based on the number of times we have sampled more extreme value. We also included raw BAF p-values from our negative control CNAs (as they were LOH variants) to have a representative null distribution.

**Matrix of Contingency based FDR estimation**

As an additional FDR check, we filled the contingency matrix of deletions and duplications, distinguishing between deletions and duplications that show positive/negative intensity in arrays. If ClinCNV called variants randomly, we could expect the array intensity to be independent of the CNA type called – array intensity may be shifted positively and negatively for false positive (or simulated) deletions or duplications with an equal probability (and it was observed for the simulated CNAs). Doubled the number of deletions with positive array intensity and duplications with negative array intensity also provides us an estimation of FDR, but for deletions/duplications only.

**The differences in FDR estimation by two methods**

The differences in FDR estimations by different methods (p-values and contingency matrix) is explained by the fact that raw normalized array values were used for contingency matrix construction and p-values from intensity and BAFs were merged for another method of FDR estimation, so, even if the breakpoints were detected inaccurately or CNV type (i.e., deletion instead of duplication or vice versa), p-value may still be quite low. We have discovered many CNAs where CNV type was incorrect in CNV-kit results and assumed that we could switch the labels of tumor and normal samples, however, all big CNAs were correctly identified by CNV-kit.

FDR for FACETS was lower with p-value estimation due to the comparatively large number of false positive deletions (non copy-neutral CNAs).

**Checking Array-based results in WES data**

Only those alterations detected in microarray data were taken into account which had more than five coverage markers in WES data or more than 5 BAF markers from WES. We took normalised coverage from ClinCNV and considered a variant as True Positive (and potentially detected in WES data) if the median Z-score of the WES coverages of the regions located within CNA borders or median Z-score of BAFs was bigger than 0.95 quantile of the normal distribution or if the absolute value of coverage was bigger than $\log_2(5/4)$ or smaller than $\log_2(3/4)$. We also looked at the direction of change in the case of copy-number imbalanced events.

#### iii.2 Recurrent CNAs in CLL

In order to investigate which genes were recurrently altered in CLL samples we used GISTIC ([Mermel et al., 2011], genomic regions signifi-

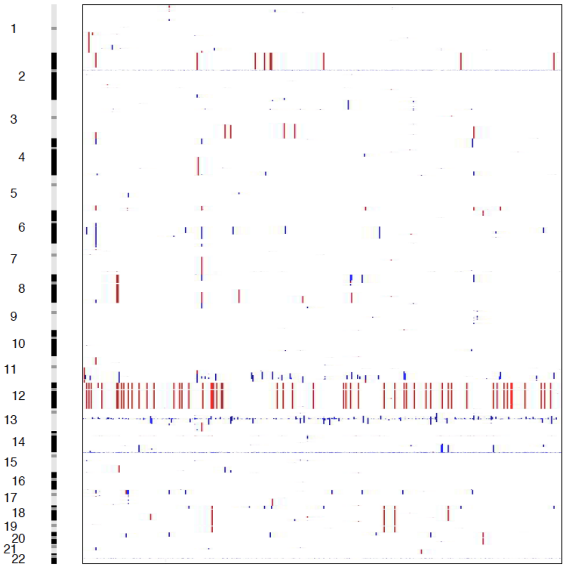


**Figure 5:** *Copy-number alterations across the whole cohort of CLL samples.*

cance plots are provded in fig. 6).

Most significant peaks occured close to centromere chr2, end of q-arm of chr14, close to centromeres of chr22 and chr14 – regions where frequently rearranged immunogenic genes are located. Other peaks contain well-known cancer genes such as TRIM13, TMPRSS5, BIRC2, TMEM123, PRSS1, SMARCC1, PARP1, etc. Whole chr12 was often duplicated.

#### iii.3 Clonal composition in CLL

Among samples that had at least one CNA detected by ClinCNV only 2 samples were detected as homogeneous. 393 samples had 2 clones, 31 samples had 3 clones and 4 samples had 4 clones, according to ClinCNV's evaluation. Frequency of clonal composition (normalised by the biggest estimated cell fraction of CNAs from corresponding samples) is depicted in the violin plot in fig. 7.

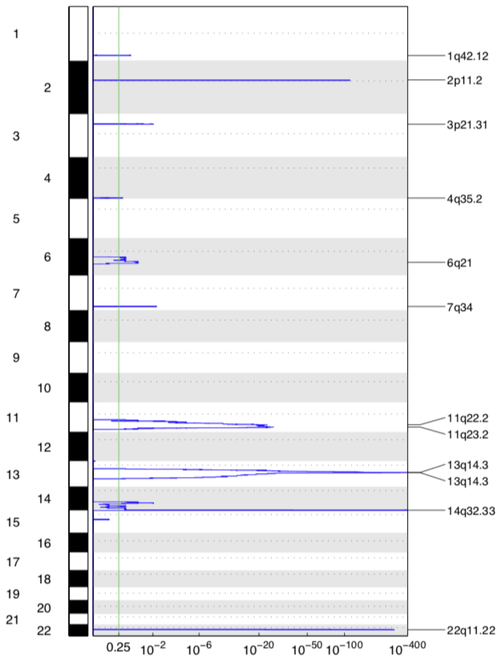


**(a)** *Enrichment of deletions analyzed with GISTIC*

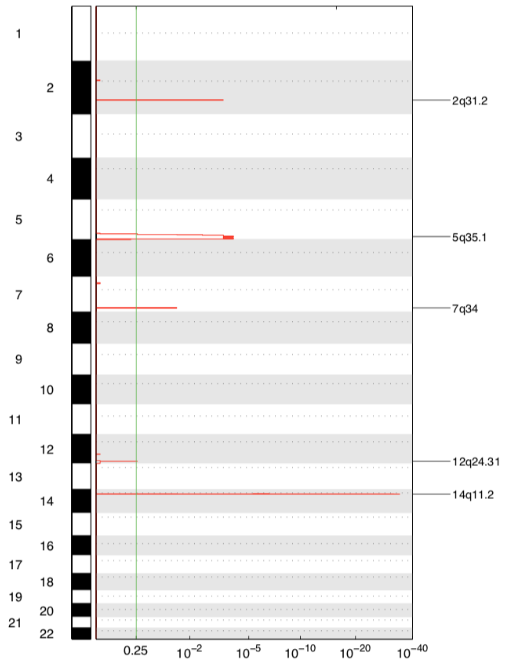


**(b)** *Enrichment of duplications analyzed with GISTIC*

**Figure 6:** *Enrichment of CNAs analyzed with GISTIC.*

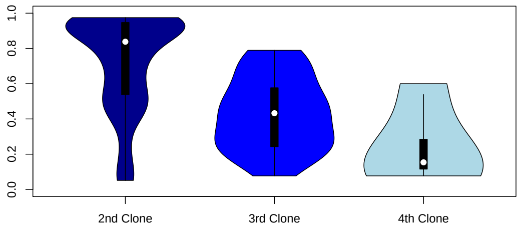


**Figure 7:** *Density of CCF of discovered clones in CLL (relatively to the biggest, 1st clone).*

### iv. In-house TPS cohort results

Speaking about the practical importance of CNAs detected, we analysed our samples with at least 1 CNA detected and with purity estimated bigger than 20% (213 samples overall) with the Cancer Genome Interpreter (CGI, [Tamborero et al., 2018]). We observed that in the majority of cases such CNAs were important: they could be annotated as druggable alterations or indicated potential resistance to particular drugs. For visualization purposes, we merged samples of cancer type that can not be directly classified into CGI categories or cancers that have less than 3 cases into one category ("Mixed"). Three bar plots (biomarkers of sensitivity, biomarkers of resistance, and biomarkers of toxicity) are provided in fig. 8.

The most commonly altered genes, annotated with the CGI, were:

TPMT (117 cases, genes play role in immune system supression, amplification leads to resistance to Cisplatin and increased toxicity to Thioguanine (Guanine analog) and Mercaptopurine (Purine analog), deletion leads to increased toxicity to above mentioned drugs according to FDA guidelines),

DPYD (89 cases), amplification or deletion leads to increased toxicity to Tegafur (Fluoropyrimidine), Flourouracil (Fluoropyrimidine), Capecitabine (Fluoropyrimidine) according to CPIC and FDA guidelines;

UGT1A1 (81 cases), alterations lead to increased toxicity to Pazopanib (VEGFR inhibitor), Irinotecan (TOPO1 inhibitor), Nilotinib (BCR-ABL inhibitor), Irinotecan (TOPO1 inhibitor), Nilotinib (BCR-ABL inhibitor) according to FDA guidelines.

#### v. Time complexity and WGS samples

ClinCNV takes approximately 3 hours for preparation of pre-calculated coverage values and BAFs for a cohort of 160 WES samples and spends from 10 minutes to one hour per WES sample analysis (depending on the number of CNAs and outliers and clonal structure complexity) using 4 cores of Intel Xeon 2.8 GHz processor, which is much slower than other tested tools, but tolerable in both research and clinical practice. Targeted panel sequenced samples are analysed for 5-10 minutes per sample using laptop with 2GHz Intel i7 processor and require around half an hour for pre-processing of around 200 samples. ClinCNV was not tested in whole genome samples, however several simplifications (such as decreasing resolution of distinct clonal fractions from 2.5% to 5%) may be applied which dramatically reduce the computational time.

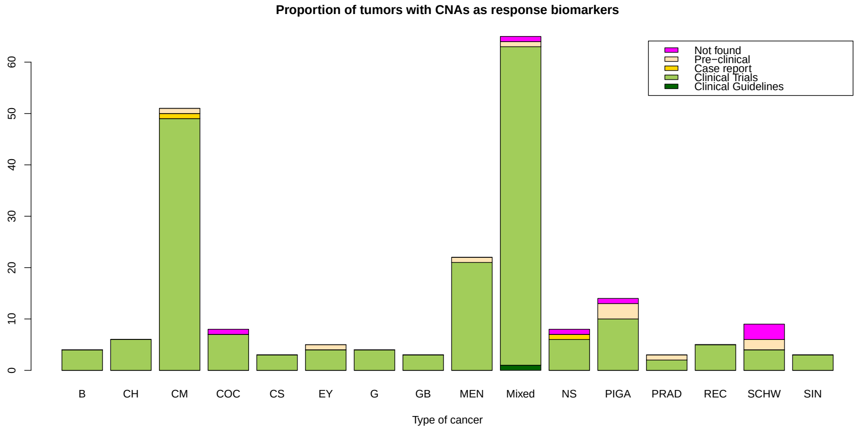


**(a)** *Statistic on genomic alterations as biomarkers of drugs response*

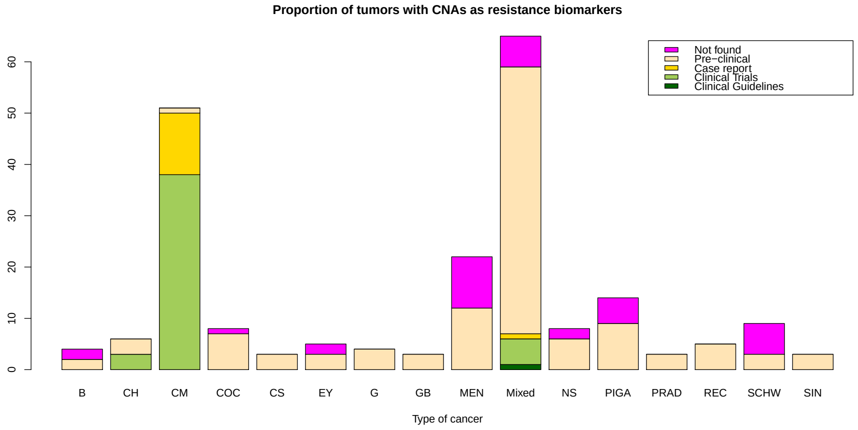


**(b)** *Statistic on genomic alterations as biomarkers of drugs resistance*

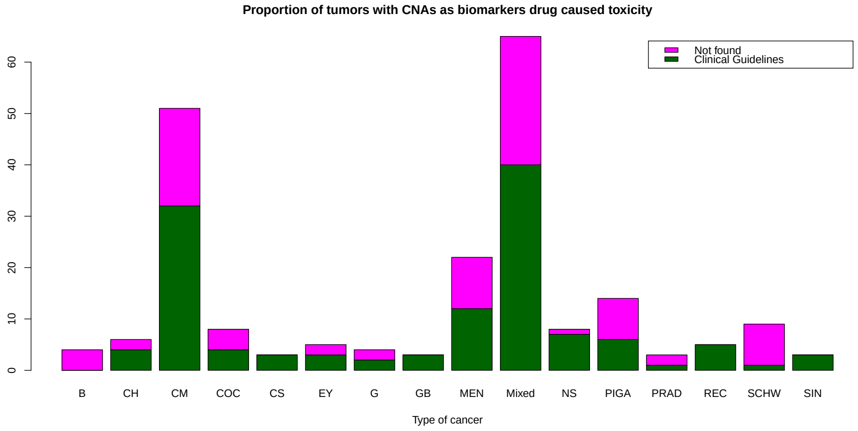


**(c)** *Statistic on genomic alterations as biomarkers of drugs' toxicity*

**Figure 8:** *Genomic alterations with the evidence of clinical impact found by ClinCNV in different tumors. Only tumors with at least one CNA detected are shown. Rare tumors from our cohort (2 or less tumors of the particular type) were not included.*
